# Novel Adult Cortical Neuron Processing and Screening method illustrates Sex- and Age-dependent effects of pharmaceutical compounds

**DOI:** 10.1101/2022.03.29.486299

**Authors:** Arthur Sefiani, Ivan Rusyn, Cédric G. Geoffroy

**Affiliations:** Department of Neuroscience and Experimental Therapeutics, College of Medicine, Texas A&M University, Bryan, TX 77807, USA; Department of Veterinary Integrative Biosciences, College of Veterinary Medicine & Biomedical Sciences, Texas A&M University, College Station, TX 77843, USA

**Keywords:** Neuroregeneration, Neuroprotection, Drug Discovery, Neuritogenesis, Assay Development, Aging, Sex Differences

## Abstract

Neurodegenerative diseases and neurotraumatic injuries are typically age-associated disorders that can reduce neuron survival, neurite outgrowth, and synaptic plasticity leading to loss of cognitive capacity, executive function, and motor control. In pursuit of reducing the loss of said neurological functions, novel compounds are sought that promote neuron viability, neuritogenesis, and/or synaptic plasticity. Current high content *in vitro* screenings typically use cells that are iPSC-derived, embryonic, or originate from post-natal tissues; however, most patients suffering from neurodegenerative diseases and neurotrauma are of middle-age and older. The chasm in maturity between the neurons used in drug screens and those in a target population is a barrier for translational success of in vitro results. It has been historically challenging to culture adult neurons let alone conduct screenings; therefore, age-appropriate drug screenings have previously not been plausible. We have modified Miltenyi’s protocol to increase neuronal yield, neuron purity, and neural viability at a reduced cost to expand our capacity to screen compounds directly in primary adult neurons. Furthermore, we have developed the first morphology-based screening system using adult cortical neurons. By using primary adult cortical neurons from mice that were 4 to 48 weeks old for screening of a library of pharmaceuticals, we have demonstrated age- and sex-dependent effects on neuritogenesis and neuron survival *in vitro*. Utilizing age- and sex-appropriate *in vitro* models to find novel compounds increasing neuron survival and neurite outgrowth, made possible by our modified adult neuron processing method, will greatly increase the relevance of in vitro screening for finding neuroprotective compounds.

## Introduction

The median age is rising worldwide^1^ as a larger percentage of the population reaches late adulthood. In the United States from 1900 to 1996, the number of people over 65 years old increased by 11-fold and the number of people over 85 years old increased by 31-fold^2^. The percent of the population over 65 doubled from 8% in 1950 to 16% in 2018^3^. Both the absolute and relative number of older individuals are rising which increases the prevalence of age-associated neurological disorders, such as neurodegenerative disorders and neurotrauma. From the year 1990 to 2010, dementia cases worldwide increased by 99.3% and the per capita rate for dementia worldwide increased by 53.3%^4^. The prevalence of traumatic brain injury (TBI) in senior citizens increased 53.5% from 2001 to 2010 as it simultaneously decreased for young adults^5^. From 1978 to 2005, the percentage of people with spinal cord injury (SCI) being of geriatric age increased by 267%^6^. Therefore, there is a growing need for developing novel therapeutic strategies specifically for protecting cognitive abilities in the older population.

In the search for novel medicines for neurodegenerative diseases and neurotrauma, researchers are looking for compounds that increase neuron survival and neurite regenerative capacity. The survival of neurons decreases after the onset of neurodegenerative diseases^7^ and neurotrauma^8,9^, escalating the pathological and neurobehavioral outcomes. Because humans have an extremely limited capacity to regenerate neurons^10^, compounds that promote neuron viability are of great interest to mitigate neuron loss and associated disease conditions. In the case of SCI, there are several neuroprotective agents undergoing clinical trials, such as Riluzole^11^, Glyburide^12^, and Minocycline^13^. Although these compounds have unique mechanisms of action, their use is intended to prevent necrosis at the injury site and mitigate further neural degeneration^14^. For Alzheimer’s disease, neuroprotective agents have been proposed to mitigate the deleterious effects of amyloid beta accumulation^15^.

Neurite growth is a vital step in the recovery process from neurotrauma to reconnect damaged connections^16^. In the case of SCI, axons extended from cortical neurons are damaged and degenerated which then must be regenerated past the lesion site to restore connection to caudal neurons^17^. Neurodegenerative diseases also induce axonal degeneration which can lead to neuronal death and progression of pathology^18^. Current therapeutic strategies in neurodegenerative diseases, such as glaucoma and Parkinson’s disease, aim at increasing axon regenerative capacity to mitigate pathology^19,20^. Aging itself reduces neurite regenerative capacity^21^ and increases susceptibility of neurons to death and degeneration^22^ which can produce worst outcomes after neurotrauma^23^ and increases the prevalence of neurodegenerative diseases^24,25^. Therefore, it is imperative that novel therapeutic strategies can increase the survival and neurite regenerative capacity of neurons in older individuals as well.

An efficient way to identify compounds that increase the survival and neurite regenerative capacity of neurons is to perform high content *in vitro* screenings. To date, such screens have involved the use of cell lines, embryonic neurons, newborn neurons, and iPSC-derived neurons^26–30^. However, these neurons do not represent the target neuron population in humans in terms of age characteristics. Indeed, aged neurons have different characteristics relative to neurons from younger individuals^21,22,31^. This dichotomy in age between *in vitro* assays and *in vivo* settings is likely to play a role in the high number of failures when moving a drug forward along the clinical phases. In fact, the age factor and associated complications have been advanced as a major component of the translational failure of promising drugs in the stroke field^32,33^ where most of the pre-clinical testing are performed in very young stroke models while human patients are older^34,35^. Additionally, the same drug may have completely opposite effects depending on age, as it is the case for methylphenidate, used to treat attention deficit-hyperactivity disorder^36^. There is an age-dependent decline in axon growth after neurotrauma^37^ demonstrating the importance of age-appropriate models in pre-clinical studies.

There are also sex-based differences in pharmacological response^38^, pharmacodynamics, and pharmacokinetics^39^; women are 50-75% more likely to experience an adverse reaction to prescription medication^40^. Because of the disparity of prevalence between sexes for certain diseases, such as the disproportionality higher SCI incidence rate in men^41^ and prevalence of stroke in young women^42^, it is vital to understand sex based efficacy of compounds before moving onto clinical stages. Screening in embryonic or early postnatal neurons cannot properly take sex into account as sex differences in motor performance^43^ and brain characteristics^44^ are more apparent after puberty. Therefore, screening compounds on adult neurons *in vitro* at different age and in both sexes should be the first step in identifying the potential of compounds to enhance neuron survival and neurite growth, increasing the potential of translation success while minimizing possible harm.

To date, culturing even young adult cortical neurons has been challenging. The few protocols existing lead to very inconsistent results, with a low yields and cell viability^45–47^. This makes it very difficult, if not impossible, to test a single drug candidate on young adult neurons, let alone to test it on older cortical neurons (>6 months). Using adult cortical neurons for an age- and sex-appropriate compounds screen *in vitro* has therefore not been conceivable yet, but it would be immensely beneficial to find demographic-appropriate drug candidates. Miltenyi recently developed a system to culture very young adult brain neurons (up to 4 weeks old)^46,48,49^. We modified this technique to culture middle- and advanced-age cortical neurons^21^. In the present report, we further adapted this protocol to screen for compounds that enhance survival and neurite outgrowth and determine the demographic the compounds have most efficacy in. Indeed, not only positive hits from embryonic or postnatal cortical neurons may not affect older neurons, but it is plausible that compounds with no effect on embryonic or postnatal neurons may in fact present beneficial effects on adult neurons only. Therefore, screens utilizing adult cortical neurons may present compounds that have been prematurely dismissed as false negatives in previous screens. Developing such a system would also allow for the examination of sex-dependent effects of compounds, an essential variable in drug development^50^. This may allow for the development of future age and sex-based personalized medicine.

Our newly developed protocol increases the number of viable neurons obtained per gram of brain tissue and significantly increases the absolute volume of brain tissue an individual can process solitarily. We designed a screening and analysis workflow to minimize human input to mitigate human biases. Through a targeted screen using our novel model, we identified compounds with age- and sex-dependent effects on neuron survival and neurite outgrowth. This clearly illustrates the need to perform future screens in different age and sex groups to 1) to determine the specific demographic the compound will have most clinical effect in; to 2) to theoretically prevent the premature dismissal of compounds with no effect in iPSC/embryonic screens yet would be advantageous to adults; and to 3) reduce the false positivity rate from iPSC/embryonic screens by efficiently vetting compounds that do not show benefit to adult neurons and therefore will likely not be efficient when testing in preclinical settings.

## Methods

### Animals

This study uses young adult and middle-aged male and female wild-type C57Bl/6 mice. The young adult group is of 4-9 weeks of age and the middle-age group is of 40-48 weeks of age. All procedures were conducted according to the protocol approved by the Institutional Review Board/Animal Ethics Committee of Texas A&M University (IACUC 2018-0324).

### Cell Culture

First, 20 μL of 50 μg/mL Poly-D-Lysine (PDL, Sigma Aldrich, A-003-M) was added onto the wells of 384-well glass bottom (Brooks Life Sciences, MGB101-1-2-LG-L) or plastic bottom (Greiner-Bio, 781091) plates and incubated inside a 5% CO2 incubator at 37 °C for 48 hours. After incubation, the wells of the plates were washed 5 times with H_2_O and set to dry overnight at room temperature and were used within 24 hours after drying. After euthanization of mice, the brains were extracted and placed in cold Hank’s balanced salt solution (HBSS) followed by microdissection of the cortex. Up to 1.25 grams of cortical tissue were placed in each gentleMACS™ C Tube (Miltenyi Biotec, # 130-093-237) which contained 5 mL of 0.3 mg/mL papain (Worthington, LS003126) diluted in HBSS. The gentleMACS™ C Tubes were placed on the gentleMACS™ Octo Dissociator with Heaters (Miltenyi Biotec, # 130-096-427) with heating cuffs attached and underwent the gentleMACS Program 37C_ABDK_01 protocol. After protocol completion, the contents of the gentleMACS™ C Tube were strained through a 70 μm cell strainer (Miltenyi Biotec, # 130-110-916) placed on top of a 15 mL conical centrifuge tube. 7 mL of cold Dulbecco’s Phosphate-Buffered Saline with glucose and pyruvate (DPBS, Thermo Fisher Scientific, 14287072) was added into each of the 15 mL conical centrifuge tubes on top of the strained cells. The 15 mL tubes were centrifuged at 300×g for 10 minutes at 4 °C before aspirating the supernatant completely. Debris removal solution was made by adding 1800 μL of Debris Removal Concentrate (Miltenyi Biotec, # 130-109-398) to 6200 μL of cold DPBS. The remaining pellet inside the 15 mL tubes was resuspended with 8 mL of Debris Removal Solution. Very slowly, 4 mL of cold DPBS was dispensed on top of the debris removal solution and cell mixture in each 15 mL tube forming a clear layer on top. The 15 mL tubes were centrifuged at 3000×g for 10 minutes at 4 °C with slow acceleration and deceleration. The top clear and middle debris layers were aspirated leaving the milky mixture beneath the debris layer untouched. 6 mL of DPBS was added onto the milky mixture and mixed gently before centrifuging at 300×g for 5 minutes at 4 °C. All supernatant was aspirated afterwards. Red Blood Cell Remover Solution was made by mixing 125 μL of Red Blood Cell Lysis Solution 10× (Miltenyi Biotec, # 130-094-183) with 1125 μL of H_2_O. The remaining pellet was resuspended in 1.25 mL of Red Blood Cell Remover Solution and incubated for 10 minutes at 4 °C before the addition of 12 mL of 0.5 % bovine serum albumin (BSA, Miltenyi Biotec, # 130-091-376) diluted in DPBS. The mixture was centrifuged at 300×g for 5 minutes at 4 °C with the supernatant aspirated completely afterwards. The remaining pellet was resuspended in 80 μL of 0.5% BSA and 20 μL of Non-Neuronal Cells Biotin-Antibody Cocktail (Miltenyi Biotec, # 130-115-389) and incubated for 5 minutes at 4 °C. Cells were washed by adding 2 mL of 0.5% BSA followed by centrifugation at 300×g for 5 minutes at 4 °C followed by aspiration of the supernatant. The remaining pellet was resuspended in 80 μL of 0.5% BSA and 20 μL of Anti-Biotin MicroBeads (Miltenyi Biotec, # 130-115-389) and incubated for 10 minutes at 4 °C. After the addition of 6 mL 0.5% BSA, the mixture was flowed through 0.5% BSA primed LS columns (Miltenyi Biotec, #130-042-401). The negative fraction containing the majority of neurons was collected and centrifuged at 300×g for 5 minutes at 4 °C and resuspended in neuron media. Unless noted otherwise, neuron media consists of MACS Neuro Media (Miltenyi Biotec, # 130-093-570), 2 mM L-alanine-L-glutamine dipeptide (Sigma-Aldrich, G8541-100ML), and 1× B-27™ Plus Supplement (ThermoFisher Scientific, A3582801). Unless noted otherwise, cells were added onto PDL coated wells and placed inside a 5% CO2 incubator set at 37 °C for the stated days *in vitro* (DIV). Unless noted otherwise, 10,000 cells were plated per well with 0.056 cm^2^ growth area. The neuron isolation, processing, and plating procedure described herein is referred to as the ‘Modified Protocol’.

### RT-qPCR for determining cell culture purity

RT-qPCR assay was replicated as previously described^21^. Briefly, Directzol RNA micro-prep columns (Zymo, R2061) are used to extract RNA from neurons directly following neuron isolation. RNA concentration was measured using the Thermo Scientific™ NanoDrop 2000. Quantabio cDNA Synthesis kit (Quanta, 95047) was used to synthesize cDNA before conducting qPCR using the Quantabio PerfeCTa^®^ SYBR^®^ Green FastMix^®^ (Quanta, 95073) on the ViiA7 Real Time PCR system (Life Technologies). The neuron enrichment in the negative fraction was calculated as previously described^21^, −ΔCT of ΔNeuN against ΔGFAP and ΔGLAST using the formula: −ΔCT = −(ΔCT NeuN – (SQRT(ΔCT GFAP^2^ + ΔCT GLAST^2^))). CT was calculated for each group based on the absolute CT per primer subtracted by the respective CT of the negative control to reduce background noise. For each group, RNA was extracted from 3 separate isolation procedures from young adult males and each sample was analyzed in triplicate. No outliers were detected nor omitted.

Primers used to identify the main cellular constituents: Neurons: MAP2 (F: 5’-CTG GAG GTG GTA ATG TGA AGA TTG; R: 5’-TCT CAG CCC CGT GAT CTA CC-3’) and NeuN (F: 5’ AAC CAG CAA CTC CACCCT TC-3’; R: 5’-CGA ATT GCC CGA ACA TTT GC-3’). Astrocytes: GFAP (F: 5’-CTA ACG ACT ATC GCC GCC AA-3’; R: 5’-CAG GAA TGG TGA TGC GGT TT-3’) and GLAST (F; 5’-CAA CGA AAC ACT TCT GGG CG-3’; R: 5’-CCA GAG GCG CAT ACC ACA TT-3’). Oligodendrocytes: Oligo2 (F; 5’-GAA CCC CGA AAG GTG TGG AT-3’; R: 5’-TTC CGA ATG TGA ATT AGA TTT GAG G-3’). β-actin: (F: 5’-CTC TGG CTC CTA GCA CCA TGA AGA-3’; R: 5’-GTA AAA CGC AGC TCA GTA ACA GTC CG-3’).

### Immunocytochemistry

Cell cultures were fixed with 4% paraformaldehyde (PFA, 15 min) after the completion of the respective experiment. After fixation, immunocytochemistry was conducted by first washing the cells with DPBS 3 times, then incubating in 5% normal horse serum for 60 min to block nonspecific binding (VWR, 102643-676). Afterwards, the cells were incubated in 1:500 TUBB3 (BioLegend, 801202) for 16 hrs. followed by another 3 washes with DPBS and incubation in 1:500 Alexa Flour 488 (ThermoFisher Scientific, A32723) and 1:10000 DAPI (VWR, 95059-474) for 60 min, all conducted at room temperature. After the completion of immunocytochemistry, the cells were preserved in Fluoromount-G Mounting Medium (ThermoFisher Scientific, 00-4958-02) until imaging.

### Analysis of neurite outgrowth and survival

Representative images in figures were imaged using 20×/63× objectives on a Zeiss Axio Observer system. For automated image acquisition, the 20× magnification lens of the ImageXpress (IXM) Micro Confocal High-Content Imaging System (Molecular Devices, San Jose, CA) was used along with the Neurite Outgrowth Analysis Module in MetaXpress^®^ 6 software (Molecular Devices) for automated image analysis, a system previously used to image and analyze changes in neuron morphology^28,51–53^. Using the 20× magnification, 16 separate images (with 10% overlap) were required to sustain >90% coverage of each well while avoiding the walls.

Three variables were quantified.

1. Valid neurons: Total number of cells in a well that are both DAPI and TUBB3 positive and with total neurite outgrowth of ≥10 μm.
2. Total neurite outgrowth: Sum of the lengths of all the neurites from a valid neuron. This is then averaged over all the valid neurons in a well.
3. Average neurite length: The total length of all the neurites from a valid neuron divided by the number of neurites and branches of that cell. This is then averaged over all the valid neurons in the well.

### Data analysis

Normally distributed data was analyzed using unpaired t-test when comparing 2 means and one-way analysis of variance (ANOVA) with Tukey’s or Dunnett’s post-hoc test when comparing >2 means. Tukey’s post-hoc test is used to compare >2 means all with one another. Dunnett’s post-hoc test is used to compare ≥ 2 means to the mean of a particular group. Two-way ANOVA was used to analyze the effects of 2 independent variables on the expected outcome. Tukey’s multiple comparisons test was used to compare the means of each cohort with the means of all other cohorts. Dunnett’s multiple comparisons test was used to compare the means of each cohort to the mean of a particular cohort. Šídák’s multiple comparisons test was used to compare the means of 2 independent groups for all cohorts. Simple liner regression was conducted to determine goodness of fit (R^2^) and to determine the correlation between 2 independent variables. Linear regression t-test was used to compare the slopes of 2 independent regression lines. Analysis and graphing were conducted using Graphpad Prism 9. All graphs represent the data mean with error bars illustrating the standard error of the mean (SEM).

### Compounds

RO48 (generously provided by Drs. Lemmon, Bixby, and Al-Ali from Miami Project to Cure Paralysis, University of Miami), (S)-H-1152 (Cayman Chemical Company, 10007653), and 7-epi Paclitaxel (Cayman Chemical Company, 20741).

## Results

### Optimization of surface coating and media supplementation

We first determined if laminin coating in addition to the PDL coated surface would improve neurite outgrowth and number of valid neurons and/or mitigate the need for extra supplementation of media with B-27™ Plus (B27^+^). A 10 μg/mL laminin coating was applied for 1 hour at 37 °C in respective wells before being washed off with Dulbecco’s Modified Eagle Medium (DMEM) prior to cell plating. Primary cortical neurons isolated from young adult male mice were cultured for 2DIV on PDL coated wells with (B27^+^) or without B27^+^ supplementation (Control), or on PDL/laminin coated plates with (B27^+^ & Laminin) or without B27^+^ supplementation (Laminin). The average neurite length (Figure 1A), total neurite outgrowth (Figure 1B), and number of valid neurons (Figure 1C) were analyzed and expressed as a percentage change relative to the control group containing no B27^+^ supplement or laminin coating. The B27^+^ supplementation induced a significant increase in average neurite length (P<0.05), total neurite outgrowth (P<0.0001), and number of valid neurons (P<0.0001). The extra laminin coating did not induce any significant changes with or without B27^+^ Plus supplementation, therefore, laminin does not improve neurite growth or number of valid neurons, nor can it replace or enhance media supplementation. Plating neurons without PDL coating resulted in few neurons adhering to the surface without substantial neurite growth (data not included). Media supplementation and surface coating is required, especially to increase the number of valid adult cortical neurons. Only PDL coating was used for subsequent experiments and all experimentation was done in standard size 384-well plates.

**Figure 1.**
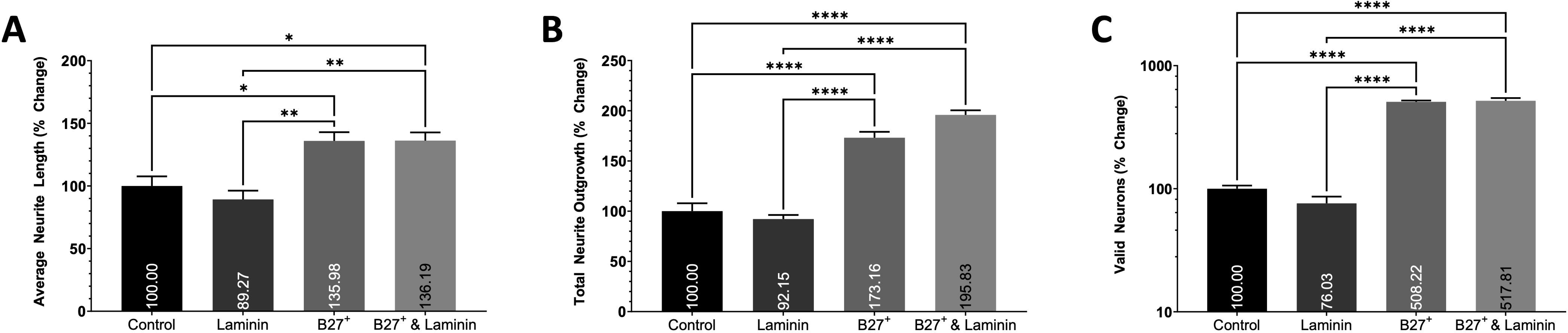
The effects of Laminin coating and media supplementation. Histograms of the A) average neurite length, B) total neurite outgrowth, and C) number of valid neurons expressed as percent change relative to the control group containing no B27+ supplement or laminin coating. Data analyzed using one-way ANOVA with Tukey’s post-hoc test comparing the mean of each condition to one another. Primary cortical neurons isolated from young adult male mice and cultured for 2DIV. 3 wells per condition. * (P<0.05), ** (P<0.01), *** (P<0.001), **** (P<0.0001). Graphs show mean and SEM.

### Determine the best digestion enzyme

Next, we determined the effectiveness of different digestion enzymes and concentrations to use to isolate cortical neurons and their efficacy to maintain their viability. Identical dissection method, dissociation method and temperature, and volumes of digestive enzymes were used herein. After the respective digestive enzymes were used in replacement of 0.3 mg/mL papain (methods section) to isolate cortical neurons from young adult male mice (1 mouse cortex per condition), the neurons were evenly distributed between 6 wells and cultured for 2DIV to measure how many cortical neurons can be extracted using each digestive enzyme and their ability to maintain neuron viability. The average neurite length, total neurite outgrowth, and number of valid neurons (Figures 2A-C) were analyzed and expressed as a percentage change relative to the MACS® P&A enzymes included in the adult brain dissociation kit. 20× magnification images were taken of the cortical neurons isolated from young adult mice using MACS P&A (Figure 2D) and 0.3 mg/mL papain (Figure 2E). Overall, papain presented beneficial effects on cell survival and neurites outgrowth compared to the MACS^®^ P&A digestive enzyme. 0.3-1.0 mg/mL papain resulted in increased total neurite outgrowth (P<0.0001) and number of valid neurons (P<0.0001) when compared to MACS® P&A. 0.1 mg/mL papain had significantly higher total neurite outgrowth and number of valid neurons compared to MACS^®^ P&A (P<0.0001), yet significantly lower values relative to higher papain concentrations (P<0.0001). This suggests digesting with papain has a dose-dependent effect that plateaus at ≤0.3 mg/mL. Notably, 0.1 mg/mL papain had higher average neurite length compared to 0.5 mg/mL papain (P<0.01) and MACS® P&A(P<0.001), although still had similar total neurite outgrowth compared to 0.5 mg/mL papain. There were no significant differences between papain concentrations of 0.3-1.0 mg/mL for any analysis, and therefore, the concentration of 0.3 mg/mL, the lowest effective papain concentration, was used for the remainder of the experiments to reduce cost and potential off-target effects induced by higher concentrations. Accutase^®^ and 0.25% Trypsin were also tested and yielded poorer outcomes compared to MACS^®^ P&A (data not included).

**Figure 2.**
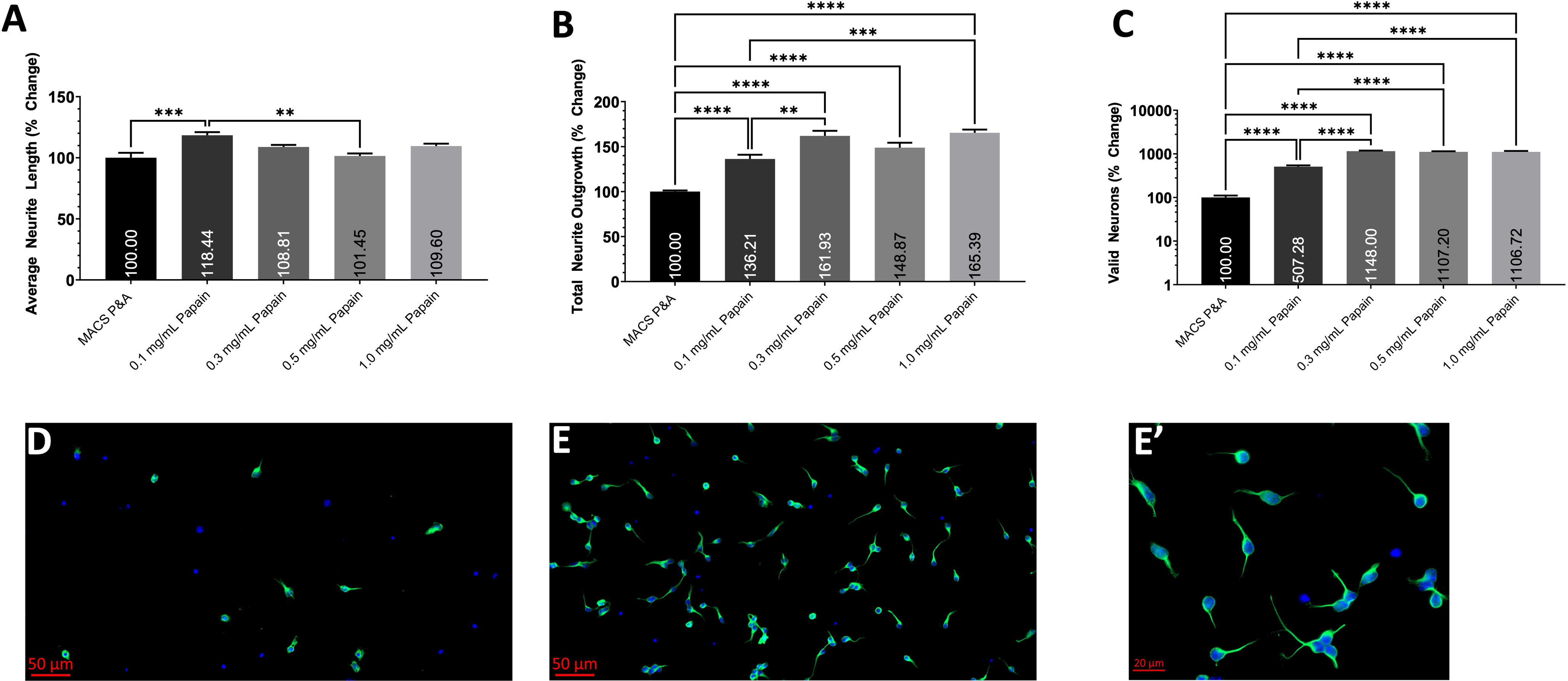
The effects of different digestion enzymes and concentrations. Histograms of the A) average neurite length, B) total neurite outgrowth, and C) number of valid neurons expressed as percent change relative to the MACS P&A group. Data analyzed using one-way ANOVA with Tukey’s post-hoc test comparing the mean of each condition to one another. Representative 20× magnification images of primary cortical neurons isolated using the D) MACS P&A and E) 0.3 mg/mL Papain (E’) 63× magnification) dissociation enzymes from young adult male mice and cultured for 2DIV; stained with TUBB3 (Green) and DAPI (Blue). 6 wells per condition. * (P<0.05), ** (P<0.01), *** (P<0.001), **** (P<0.0001). Graphs show mean and SEM. Scale Bar = 50 μm (D, E) or 20 μm (E’).

### Determine the best dissociation method, timing, and temperature

Here, we determined the effectiveness of different digestion methods, the incubation timing of those methods, and the incubation temperatures on the dissociation of cortical tissue from young adult male mice. After cortical neurons are extracted with each respective protocol, the cells are evenly dispersed between 6 wells to analyze both the yield and viability of the extracted cortical neurons. The average neurite length, total neurite outgrowth, and number of valid neurons are expressed as a percentage change relative to the ABDK (30 min) group. The gentleMACS Program 37C_ABDK_01 protocol, referred to as ABDK (30 min), is the standard protocol conducted at 37 °C using the gentleMACS™ Octo Dissociator with Heaters as per Miltenyi instructional manual. To reduce the stress on cortical neurons, the duration of the protocol was shortened to 10 and 20 min. To test the need and efficacy of the gentleMACS™ Octo Dissociator with Heaters, similar conditions were created by installing a revolving apparatus in the Thermo Scientific™ MaxQ™ 8000 Incubated Stackable Shakers that induced shaking of the contents inside. Neurons dissociated using the gentleMACS™ Octo Dissociator with Heaters had longer average neurite lengths and total neurite outgrowth in comparison to neurons incubated in a revolving apparatus. The incubation time had no effect on neuron morphology when using gentleMACS™ Octo Dissociator with Heaters, yet, when using a rotating apparatus, a reduction in time lead to reduced total neurite growth (P<0.0001). The number of valid neurons was affected by method, timing, and temperature. Using the gentleMACS™ Octo Dissociator with Heaters increased the number of valid neurons by approximately 4-fold regardless of the incubation time relative to using a revolving apparatus at 37 °C (P<0.0001). In each dissociation method, there was approximately a 2-fold increase in the number of valid neurons for every 10 min increase in incubation time (P<0.0001). Furthermore, reducing the temperature from 37 °C to 25 °C during the 20 min incubation in a revolving apparatus reduced the number of valid neurons without impacting neuron morphology (P<0.0001, Figure 3C). The gentleMACS Program 37C_ABDK_01 protocol was used in the following experiments.

**Figure 3.**
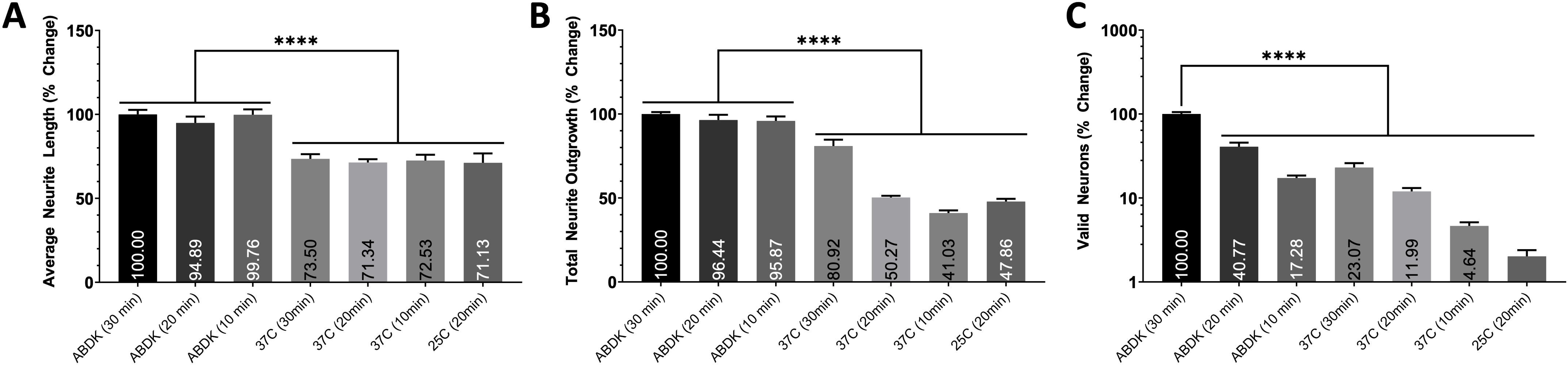
The effects of different dissociation protocols, timing, and temperature. Histograms of the A) average neurite length, B) total neurite outgrowth, and C) number of valid neurons expressed as percent change relative to the standard gentleMACS Program 37C_ABDK_01 protocol (ABDK (30 min)). Data analyzed using one-way ANOVA with Dunnett’s post-hoc test comparing the mean of each condition to the mean of ABDK (30 min). In parentheses represents the length of the protocol whether being ran on the gentleMACS™ Octo Dissociator with Heaters (ABDK) or in a revolving apparatus incubated at 37 °C or 25 °C. Primary cortical neurons isolated from young adult male mice and cultured for 2DIV. 6 wells per condition. * (P<0.05), ** (P<0.01), *** (P<0.001), **** (P<0.0001). Graphs show mean and SEM.

### The effects of cell plating density

The reaction of neurons to compounds may be dependent on their plating density, therefore, we assessed the effects of cell plating density of young adult cortical neurons on the average neurite length, total neurite outgrowth, and number of valid neurons per well (Figure 4). Cortical neurons were plated at 1,500, 3,750, 5,000, 7,500, 10,000, and 15,000 cells per well, in presence of (S)-H-1152 or Vehicle. (S)-H-1152 is a selective and potent rho-associated kinase (ROCK) inhibitor that attenuates KCl-induced contractions of femoral arteries^54^ and augments neurite outgrowth in dorsal root ganglion cells isolated from 1-day old rats that are cocultured with Schwan cells^55,56^. A simple linear regression analysis showed there was no significant correlation between plating density and the average neurite outgrowth for both 0.05% DMSO (Vehicle) and 5 μM (S)-H-1152 treated groups. There was a significant positive correlation between both total neurite outgrowth (P values: Vehicle=0.0006, (S)-H-1152=0.0001) and number of valid neurons per well (P values: Vehicle=0.0006, (S)-H-1152<0.0001) and the cell plating density for both treatment groups. Only the slope of the total neurite outgrowth linear regression line differed significantly between treatment groups, (S)-H-1152 induced a steeper increase in total neurite outgrowth and therefore more responsive to increase plating density (P<0.05). Using two-way ANOVA to compare the means of different treatment groups at each respective plating density, (S)-H-1152 significantly increased the average neurite length (P<0.05) and number of valid neurons (P<0.01) only when 10,000 cells were plated per well. Therefore, plating density can determine which compounds are deemed positive hits and must be taken into consideration during screenings. A quadratic least square regression determined the number of valid neurons for both Vehicle (R^2^=0.9911) and (S)-H-1152 (R^2^=0.9923) increases exponentially, therefore, there may be extrinsic factors related to plating density that augment the neurite formation and/or survival of neurons. To minimize the impact of these factors when screening for new compounds, which may hinder the effect of the tested drugs, the concentration of 10,000 neurons per well appears the best choice and was used in the rest of the experiments.

**Figure 4.**
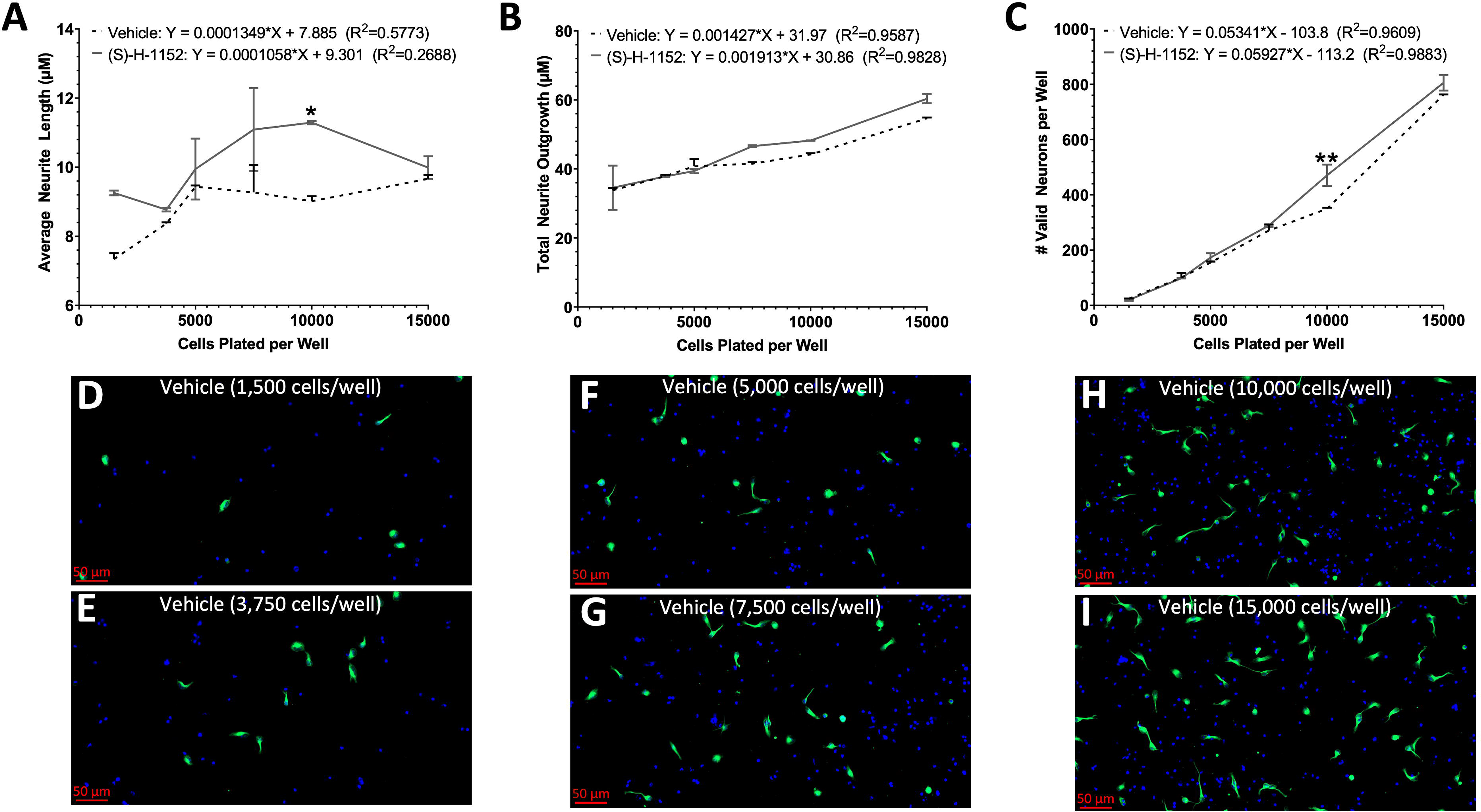
The effects of cell plating density. Linear trends of the A) average neurite length, B) total neurite outgrowth, and C) number of valid neurons of Vehicle and (S)-H-1152 treated primary cortical neurons isolated from young adult male mice cultured for 2DIV. The X-axis denotes the number of cells plated in each well. Representative 20× magnification images of primary cortical neurons from young adult male mice treated with Vehicle for 2DIV and plated at D) 1,500 cells/well, E) 3,750 cells/well, F) 5,000 cells/well, G) 7,500 cells/well, H) 10,000 cells/well, and I)15,000 cells/well; stained with TUBB3 (Green) and DAPI (Blue). Two-way ANOVA with Šídák’s multiple comparisons test was used to compare the means of neurons with different treatments for each respective plating density denoted by * (P<0.05), ** (P<0.01), *** (P<0.001), **** (P<0.0001). Simple liner regression conducted to determine goodness of fit and if the slope differs significantly from 0. Linear regression t-test was used to compare the slope of the regression lines. 2 wells per condition. Graphs show mean and SEM. Scale Bar = 50 μm.

**Figure 5.**
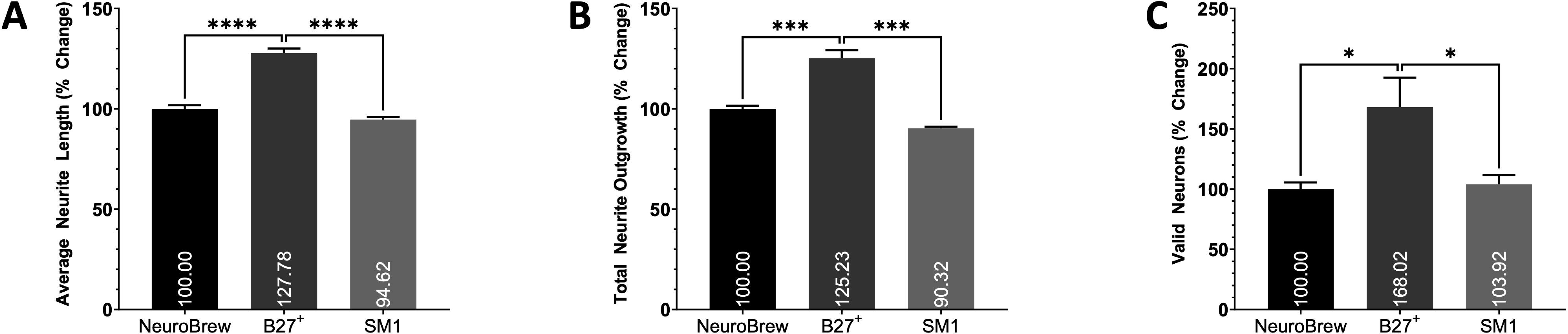
The efficacy of various neuronal supplements. Histograms of the A) average neurite length, B) total neurite outgrowth, and C) number of valid neurons of primary cortical neurons treated with the respective neuronal supplement isolated from young adult male mice and cultured for 2DIV. Data is expressed as percent change relative to the B27+ supplement cohort. Data analyzed using one-way ANOVA with Dunnett’s post-hoc test comparing the mean of each condition to the mean of the B27+ supplement group. 3 wells per condition. * (P<0.05), ** (P<0.01), *** (P<0.001), **** (P<0.0001). Graphs show mean and SEM.

### The efficacy of various neuronal supplements

To improve the survival of neurons and create an environment that resembles *in vivo* conditions, neuron supplements are added to media to study the synaptic function, neurite growth, and survival of primary neurons *in vitro* in a chemically defined manner without the use of serum^57–60^. Here, we determined the efficacy of MACS® NeuroBrew®-21 (NeuroBrew, Miltenyi Biotec), B27^+^ (Gibco), and NeuroCult™ SM1 (SM1, Stemcell Technologies) as serum-free neuronal supplements to support neuron survival and neurite outgrowth. Primary cortical neurons from young adult male mice were plated at 10,000 cells per well and the effects of the different supplements on the average neurite length, total neurite outgrowth, and number of valid neurons were assessed as a percentage change relative to the NeuroBrew group. All supplements were used according to their respective instructional manuals and added to MACS® Neuro Medium. The use of B27^+^ resulted in an increase in the average neurite length (P<0.0001), total neurite outgrowth (P<0.001), and number of valid neurons (P<0.05) relative to all other cohorts. Therefore, B27^+^ will be used in future screenings to better nurture the adult neurons. The potential issue in using B27^+^ is the fact that it may mask the true effects of compounds due to its increased potency compared to other supplements, although, compound screenings cannot be done in neurons that are not viable and reactive to drug treatment.

### The effects of the vehicle DMSO

Due to the ability of dimethyl sulfoxide (DMSO) to dissolve hydrophobic compounds^61^, it is commonly used as a vehicle in high content screenings^62,63^. To determine the tolerance of adult neurons to DMSO, we plated primary cortical neurons from young adult male mice in presence of different concentrations of DMSO and analyzed average neurite length, total neurite outgrowth, and number of valid neurons (Figure 6), expressed as a percentage change relative to the negative control (0% DMSO). The linear regression analysis confirmed a significant negative correlation between average neurite length (P=0.0404), total neurite outgrowth (P=0.0019), and number of valid neurons per well (P=0.0055) and the percentage of the media containing DMSO (% DMSO). Therefore, DMSO concentrations must be limited to the absolute minimum possible (≤ 0.075%) to avoid the harmful effects on neurons.

**Figure 6.**
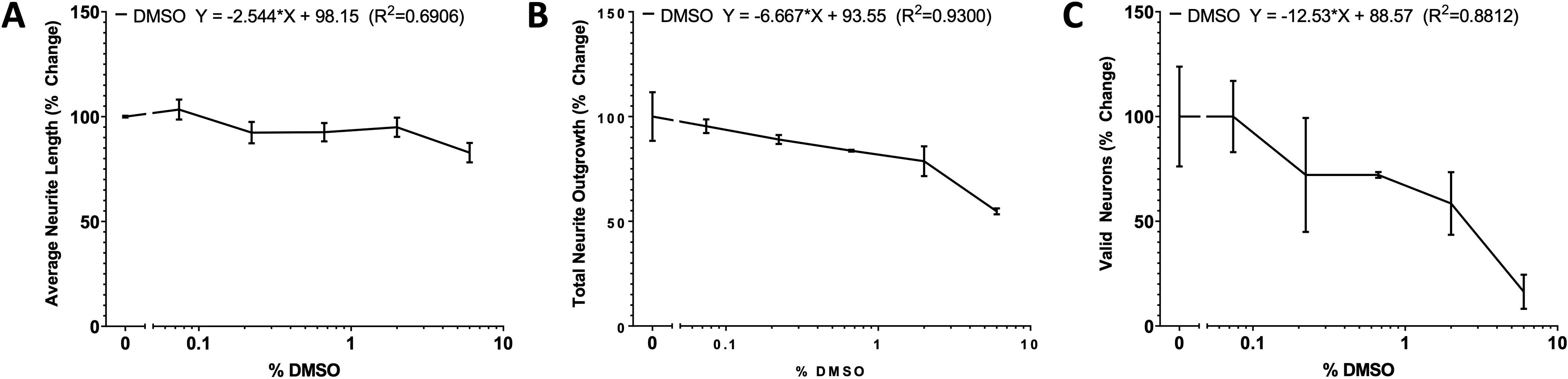
The effects of the drug vehicle DMSO. Linear trends of the A) average neurite length, B) total neurite outgrowth, and C) number of valid neurons of primary cortical neurons isolated from young adult male mice cultured in various percentages of DMSO for 2DIV. The X-axis denotes the percentage of the neuron media comprising of DMSO. The values are expressed as percent change relative to the 0% DMSO cohort. Simple liner regression conducted to determine goodness of fit and if the slope differs significantly from 0. 2 wells per condition. Graphs show mean and SEM.

### Culture purity assessed with RNA expression analysis

The differences between the original and modified neuron isolation, processing, and culturing protocols are evident in the analysis for neurite outgrowth and valid neurons. These modifications have allowed for the culturing and screening of older adult cortical neurons. We performed RNA expression analysis using RT-qPCR to determine how these modifications have affected the culture purity, by assessing RNA yield, neuron specific yield, and purity (Figure 7). The cells isolated from cortical tissue of young adult male mice were pelleted after the completion of the respective protocol followed by the extraction of RNA immediately after. The RT-qPCR data indicates that the modified protocol resulted in significantly more RNA being isolated from each cortex (P<0.0001), greater NeuN expression from the isolated cells (P<0.01), and increased NeuN expression relative to GFAP and GLAST (P<0.05). The *in vitro* cultures were suggestive of the modified protocol containing more cells; this data confirms that not only are more cells isolated, but a larger percentage of those cells are neurons. The modified protocol increased RNA yield by only 66% (Figure 7A) yet increased the NeuN expression by 112% (Figure 7B), suggesting an unproportionally larger increase in neurons relative to other neural cells. This was confirmed by the significant increase in the relative NeuN expression in the modified protocol compared to the combination of GLAST and GFAP expression (Figure 7C).

**Figure 7.**
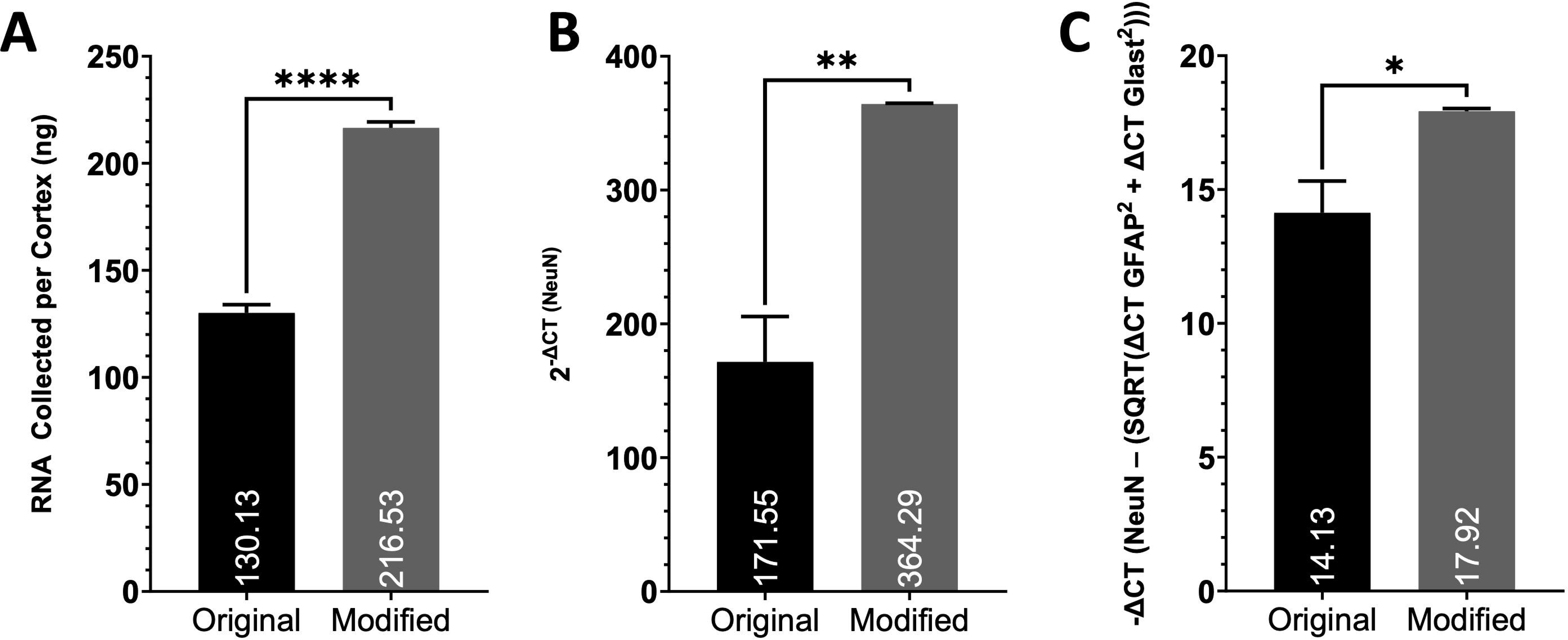
Culture purity assessed with RNA expression analysis. Histograms of the A) mass of RNA collected from each preparation, B) the relative expression of NeuN expressed as 2-ΔCT (NeuN) and C) the relative expression of NeuN in relation to GFAP and Glast comparing the original protocol from Miltenyi (Original) and our finalized modified protocol (Modified). −ΔCT = −(ΔCT NeuN – (SQRT(ΔCT GFAP2 + ΔCT Glast2))). All ΔCT values are calculated as follows: ΔCT Primer = CT Primer (Sample) – CT Primer (Negative Control). Student’s T-test was used to compare the means of each cohort. RNA extracted from primary cortical neurons isolated from young adult male mice following the original and modified protocol. 3 independent samples per cohort, each analyzed in triplicate. * (P<0.05), ** (P<0.01), *** (P<0.001), **** (P<0.0001). Graphs show mean and SEM.

### Sex and age-dependent effects of RO48 and culturing methods

Using one-way ANOVA with Tukey’s post-hoc test, we analyzed the effects of our isolation method, media, culturing protocol, and Vehicle against the young adult and middle-aged female cohorts. The middle-aged female cohort had a small but significant increase in average neurite length and significant decrease in total neurite outgrowth compared to the younger female cohort (Figure 8A-B). The middle-aged female cohort has a non-significant downward trend in number of valid neurons compared to the younger female cohort (Figure 8C). RO48 activates mammalian target of rapamycin complex (mTORC)1/2 and phosphatidylinositol-3-kinase (PI3K) and decreases the phosphorylation of S6-926^64^. To determine the sex and age-dependent effects of RO48, RO48 was added to the media of adult neurons isolated from the cortex of young adult male, young adult female, and middle-aged female mice for 2DIV and average neurite length, total neurite outgrowth, and number of valid neurons were quantified (Figure 8). 20× magnification images were taken of primary cortical neurons isolated from middle-aged female mice treated with Vehicle (Figure 8D) and 3400 nM RO48 (Figure 8E) for 2DIV. A simple linear regression concluded a significant positive correlation between the RO48 concentration in the media and the total neurite outgrowth (P=0.0019) and number of valid neurons (P=0.0335) in the young adult female cohort only. Using a Šídák’s multiple comparisons test, the means of each tested concentration of RO48 was compared to the Vehicle treated group for each age cohort and to compare the relative percent change in value 3400 nM RO48 induced between the 3 age cohorts. The average neurite length (Figure 8D) increased more, relative to the Vehicle, for the younger male and female cohorts compared to the middle-aged female cohort. The middle-aged female cohort was the only cohort without a significant increase in average neurite length at any concentration, resulting in a significant difference in the relative increase of neurite length at 3400 nM RO48 between the younger cohorts. For the total neurite outgrowth (Figure 8E), the middle-aged female cohort has a significant increase in the neurite outgrowth compared to Vehicle treatment and significantly larger relative change compared to the younger cohorts. The young adult male and female cohorts do not differ significantly, although, the young adult female cohort does have a significant increase in neurite length at the highest dose. When analyzing the number of valid neurons (Figure 8F), all 3 cohorts have a significant increase in valid neurons at doses ≥ 300 nM, although, the relative increase from the Vehicle treatment differs significantly between all 3 cohorts. The young adult female cohort is most responsive, followed by the young adult male and middle-aged female cohort. This data elucidates the sex and age-dependent effects of RO48 on adult neurons *in vitro* which are more profound at higher RO48 concentrations. More importantly, it demonstrates the need to test compounds in both sexes and age groups to determine demographic specific efficacies.

**Figure 8.**
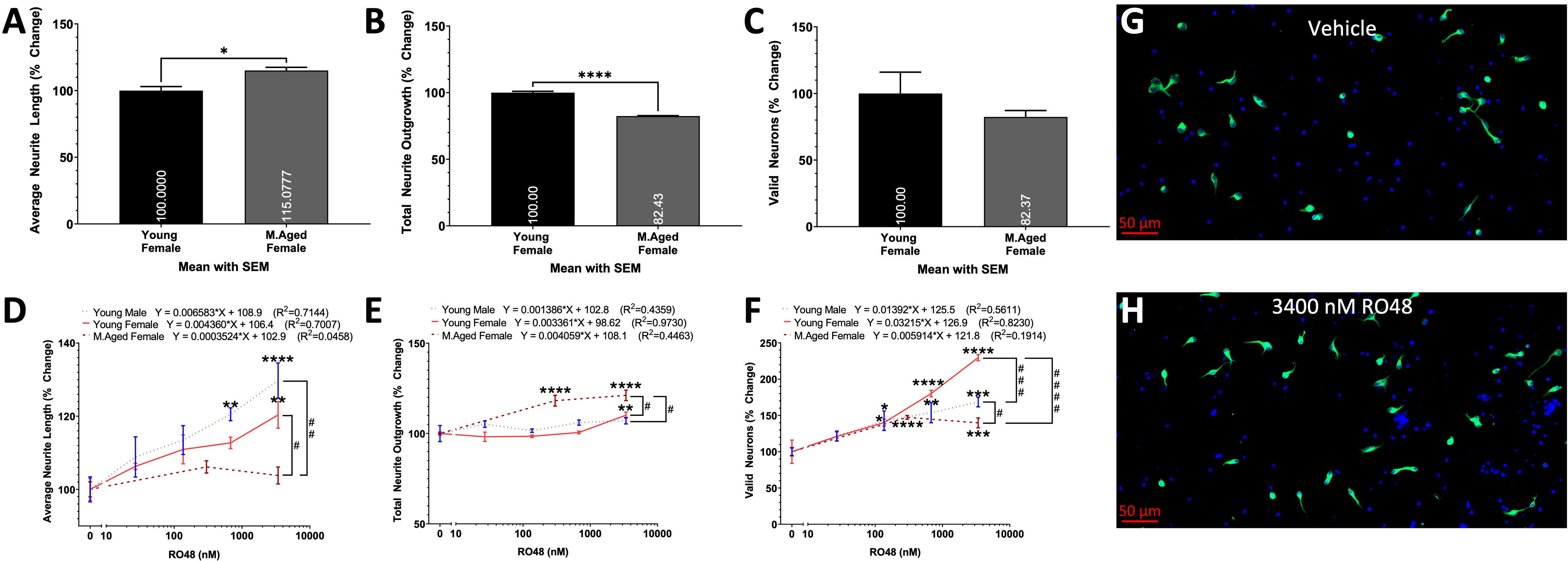
Sex and age-dependent effects of RO48. Histograms of the A) average neurite length, B) total neurite outgrowth, and C) number of valid neurons extracted from the Vehicle treated group from figures A, B, and C, respectively, expressed as percent change relative to the young adult female (Young Female) cohort. Data analyzed using student’s T-test comparing the means of each cohort denoted by * (P<0.05), ** (P<0.01), *** (P<0.001), **** (P<0.0001). Linear trends of the D) average neurite length, E) total neurite outgrowth, and F) number of valid neurons of primary cortical neurons isolated from young adult male, young adult female, and middle-aged female mice cultured in various concentrations of RO48 for 2DIV. The X-axis denotes the concentration (nM) of RO48 in the media (D, E, F). The values are expressed as percent change relative to the Vehicle treatment group of each cohort. Representative 20× magnification images of G) Vehicle and H) 3400 nM RO48 treated primary cortical neurons isolated from middle-aged female mice and cultured for 2DIV; stained with TUBB3 (Green) and DAPI (Blue). Simple liner regression was conducted to determine goodness of fit and if the slope differs significantly from 0. Linear regression t-test was used to compare the slope of the regression lines. Two-way ANOVA with Šídák’s multiple comparisons test was used to compare the means of neurons treated with 3400 nM RO48 between cohorts denoted by # (P<0.05), ## (P<0.01), ### (P<0.001), #### (P<0.0001) and to compare the means of neurons treated with various concentrations of RO48 to the Vehicle treatment group of the respective cohort denoted by * (P<0.05), ** (P<0.01), *** (P<0.001), **** (P<0.0001). All cohorts have equal parts Vehicle in media (0.05% DMSO). 3 wells per condition. Graphs show mean and SEM. Scale Bar = 50 μm.

### The age-dependent toxicity of 7-epi Paclitaxel

7-epi Paclitaxel is an FDA-approved drug for use in patients with ovarian cancer^65^. 7-epi Paclitaxel stabilizes microtubule bundles, impairs organelle transport^66^, induces peripheral neuropathy through the CXCR1/2 pathway^67^, and reduces brain injury after repeated traumatic brain injuries in mice by inducing neurite growth and nerve regeneration^68^. Adult neurons isolated from the cortex of young adult male, young adult female, and middle-aged male mice were treated with 150 nM 7-epi Paclitaxel, 3000 nM RO48, and Vehicle for 3DIV to analyze the average neurite length, total neurite outgrowth, and number of valid neurons (Figure 9). One-way ANOVA with Dunnett’s multiple comparisons test was used to compare the means of each compound to the mean of the Vehicle treated group within each respective cohort. 7-epi Paclitaxel significantly increases the average neurite length of all 3 age cohorts, yet only significantly decreases the total neurite outgrowth and number of valid neurons of the young adult cohorts without affecting the middle-aged male cohort. Therefore, screening 7-epi Paclitaxel in young adult neurons at 3DIV would yield an overall negative result and would have been prematurely dismissed while yielding an overall positive result in middle-aged male neurons. Studying the influence of 7-epi Paclitaxel in different demographics uncovers the need to conduct age-appropriate screenings to mitigate premature dismissal of potentially beneficial compounds.

**Figure 9.**
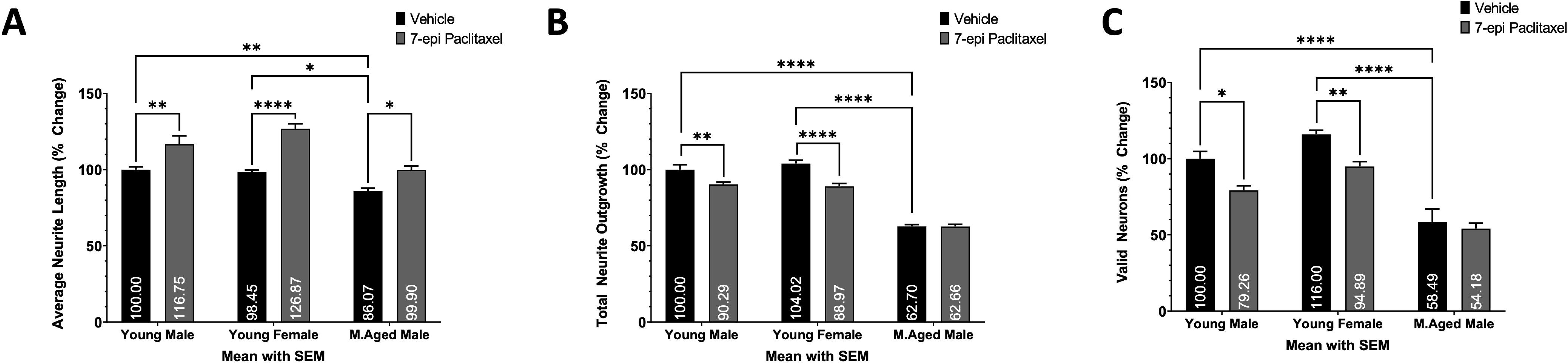
The age-dependent effects of 7-epi Paclitaxel. Histograms of the A) average neurite length, B) total neurite outgrowth, and C) number of valid neurons expressed as percent change relative to the young adult male cohort with Vehicle treatment. Primary cortical neurons isolated from young adult male, young adult female, and middle-aged male mice cultured in Vehicle or 150 nM 7-epi Paclitaxel or 3DIV. Two-way ANOVA with Dunnett’s multiple comparisons test was used to compare the means of each compound to the mean of the Vehicle treated group within each respective cohort. Two-way ANOVA with Tukey’s multiple comparisons test was used to compare the mean of the Vehicle treated groups between different age cohorts. All cohorts have equal parts Vehicle in media (0.05% DMSO). 4 wells per condition. * (P<0.05), ** (P<0.01), *** (P<0.001), **** (P<0.0001). Graphs show mean and SEM.

**Figure 10.**
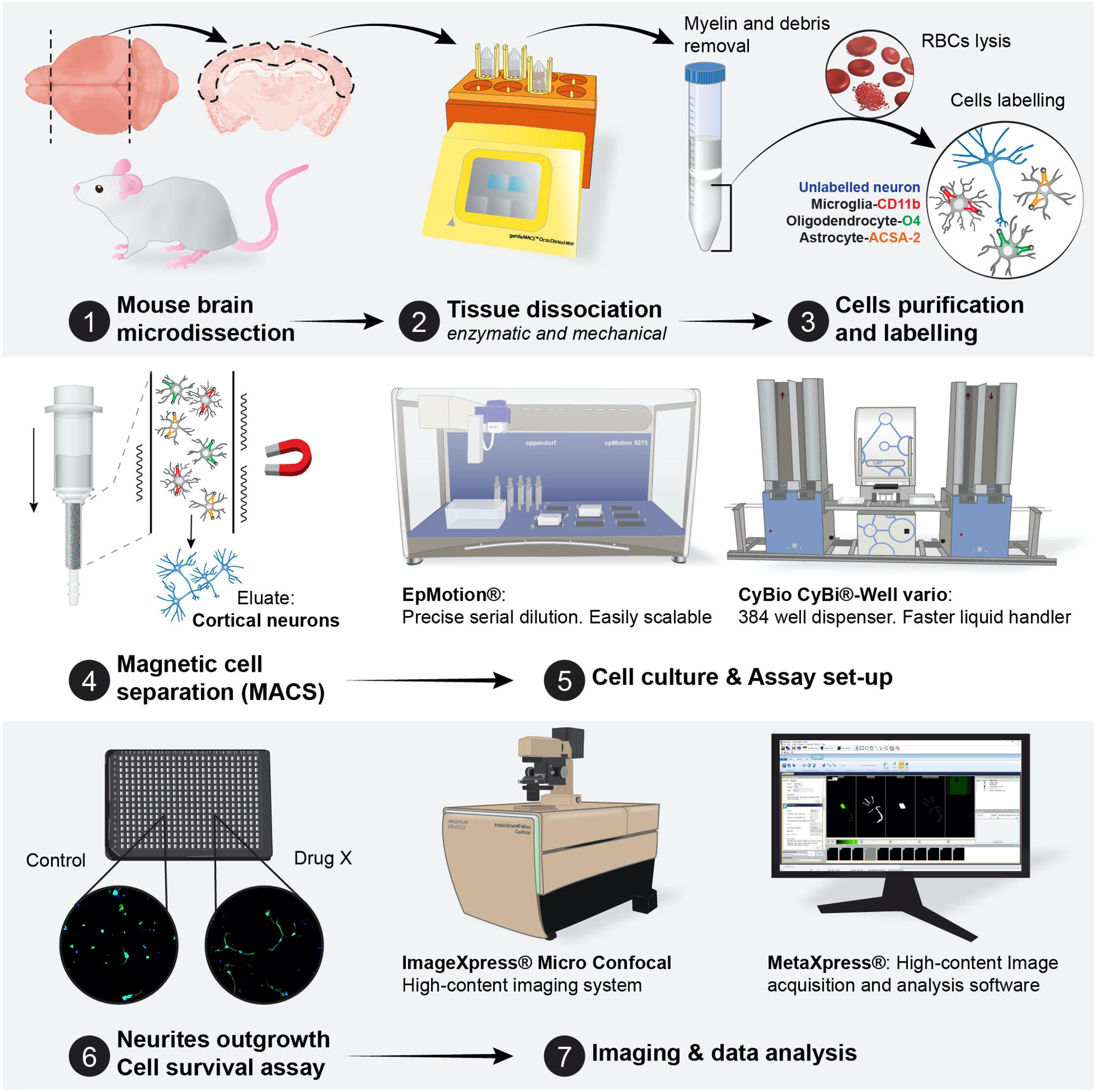
The screening platform using primary adult neural cells. Briefly, the cortex is extracted from mice, dissociated in a dissociator, followed by the removal of debris and red blood cells. Afterwards, the glial cells are separated from neurons. All liquid handling, imaging, and analysis is done with bias and human free robots.

## Discussion

We developed a new screening platform utilizing primary adult mouse cortical neurons and demonstrated sex- and age-dependent effects of neuroactive compounds. This discrepancy between demographics supports the notion that future screenings must include both sexes and different age groups to account for sex- and age-dependent processes that may alter drug efficacy and even elicit opposite effects. Therefore, screening in multiple demographics is a necessary step in reducing unforeseen erroneous results and increasing confidence in the results.

High content screenings are a vital process in drug discovery to rapidly identify potential candidates and pathways to achieve the desired therapeutic benefits. To increase the likelihood of translating said findings into a clinical setting, the assay must use cells that are comparable to the cellular targets *in vivo*. For this reason, cortical neuron screens are commonly used for finding novel therapeutics to increase axon regenerative capacity for neurotraumatic injuries^69,70^. Considering the many epigenetic^71,72^ and metabolic^21^ changes occurring in neurons as they age, the age-factor must be considered during screenings. For example, pro-inflammatory cytokines deemed harmful to adults are beneficial for the development of young neurons^25^. Therefore, compounds that modulate cytokine production are expected to produce age dependent effects. 7-epi Paclitaxel is an example of such a phenomenon for the need to use demographic specific screens. Nonetheless, screening adult, even young adult, neurons has previously not been plausible. Thanks to the advancements made by Miltenyi in neuron dissociation and isolation, small volumes of cortical neurons can be cultured. We modified this protocol to mass process larger volumes of tissue with improved methods that increase the yield and purity to conduct screenings of neurons of all ages given enough brain tissue matter.

The MetaXpress® 6 automated analysis system from Molecular Devices provides an unbiased analysis at an unprecedented rate allowing for the high content morphology-based neuron screens. The analysis used herein is based on 3 parameters: average neurite length, total neurite outgrowth, and number of valid neurons. First, the software finds nuclei (DAPI) that are associated with neurites (TUBB3). Next, the software determines if the total associated nerites are ≥ 10 μm in length and only then considers the cell to be a valid neuron. This analysis method is biased against neurons of very little neurite growth but simultaneously protects the users from falsely accounting debris particles as valid neurons. Notably, using the valid neurons analysis does not allow for the determination of how many neurons survived or how many neurons initiated neurite growth, rather analyzes a combination of the two. Therefore, with this analysis method, we cannot directly conclude that increased valid neurons increases the survival rate, although is hypothesized to be highly associated with the average viability of the entire culture. Next, the software quantifies neurite outgrowth only from these valid neurons and divides that value from the number of valid neurons to determine the average total neurite outgrowth per neuron. To determine the average neurite length, the total neurite outgrowth value is divided by the average number of branches found on only valid neurons. Therefore, the average neurite length analysis cannot determine if the neurites have equal sized branches or very long neurites with minuscule branching neurites. The software cannot specifically analyze the length of the longest neurite which is one factor in determining average neurite length. 7-epi Paclitaxel induces a noticeable increase in the length of the longest neurite (data not included) and significantly increases the average neurite length, therefore, the two analyses can be associated. More laborious analysis methods, such as neurite tracing^21^, can be used to further evaluate morphological changes. Regardless, our analyses provide enough information for determining which compounds have potential in preclinical settings and should be analyzed in further detail before further investment in preclinical trials.

Many variables went into consideration in the formation of the final neuron isolation, processing, and culturing protocol. Deciding the precise details or even inclusion of each step was determined by the effects of that step on each of the variables analyzed and the associated cost, labor, and time requirements. To assess the effects of compounds on neuron morphology, the neuron must be able to survive in culture which requires attaching to the surface and initiating neurite outgrowth and be healthy/viable enough to respond to environmental changes such as the addition of a drug. A protocol that maximizes neuron survival and viability is more versatile to other species, age groups, and regions of the CNS. Our protocol is tailored to cortical neurons, although, with slight modifications, can be used for neurons from other regions. We were able to culture young adult hippocampal and spinal neurons by adding the SM1 supplement in addition to B27^+^ and 10 ng/mL recombinant human brain derived neurotrophic factor (BDNF, Tonbo Biosciences, 21-8366-U010) to the neuron media. BDNF artificially added into culture was expected to increase the viability of neurons plated at low densities, although was contrary to our findings. BDNF has been found to stop enhancing the survival of hippocampal neurons at ages above E17^73^ even though tyrosine receptor kinase B (TrkB), the high-affinity receptor for BDNF, is present in the cortex of adult rodents^74^. It is plausible that B27^+^ contains factors that activate TrkB, therefore, the addition of more BDNF has residual effect. Increasing the concentration of debris removal solution from the recommended 22.5% to ≥25% reduces the number of valid neurons tremendously. Conversely, tissue samples containing large amounts of debris prevent the visualization and quantification of neurons and neurites. Although the removal of debris can enhance neuron survival and neurite outgrowth^75,76^, the debris removal solution may remove essential factors and biological agents that support survival and neuritogenesis. In fact, we observed improvements in survival when removing the debris removal step (not shown).

However, this is accompanied with a lot of debris that make semi-automatic quantifications difficult. Since the purpose of the assay is to evaluate increases in neurite outgrowth, the protocol is designed to ensure minimal baseline increase in neurite outgrowth without affecting the number of valid neurons. To reduce potential off-target effects, cost, and time, the protocol minimizes reagent usage. Laminin coating was deemed unnecessary and excluded from the protocol (Figure 1) and the lowest concentration of papain with maximal effect is recommended (Figure 2). In terms of deciding between neural supplements, B27^+^ significantly improves all 3 variables compared to SM1 and NeuroBrew. The potential problem with using the best supplement such as B27^+^ is it may mask the true effects of compounds by providing an overabundance of growth factors and nutrients. This theory was disproven by the significant effects seen with different drug treatments in culture supplemented with B27^+^ (Figures 8 & 9). Increasing the number of neurons through proper media supplementation allows for larger screens while reducing the variability of the analyses as there are more neurons to quantify. B27^+^ provides a nice compromise between survival and neurite growth, with a well characterized fetal bovine serum (FBS) free composition, allowing for accurate assessment of the effects of compounds on adult neurons.

The cell plating density is an important aspect of cell cultures^77^ and can dictate the survival of neurons^45,78^. For hippocampal neurons, cell plating density influences synapse formation^79^, maturation, and intensity of electrical activity^80^. Our data demonstrates the effects of plating density on viability and morphology. At densities < 5,000 cells/well, the neurons are less likely to produce the typical elongated neurites characteristic of a healthy neuron and typically too few neurons to accurately determining differences between treatment groups; variability between treatment groups is inversely proportional to the number of neurons quantified. Adult neurons plated at ≥ 15,000 cells/well have extended and overlapping neurite outgrowth that makes accurate quantification difficult. Having a plating density that is too high (≥ 15,000 cells/well) also leads to an overpopulated culture that grows at an expedited rate which can mask the effects of compounds. Therefore, subsequent experiments and the preferred plating density is 10,000 cells for a well with an area of 0.056 cm^2^. As we increased plating density, we observed an exponential increase in number of valid neurons and there is a significant correlation between neurite outgrowth and plating density. Therefore, extrinsic factors associated with plating density may influence both the number of valid neurons and neurite morphology. One potential factor is glutamine released by neighboring neurons^81^. Glutamine is artificially added to the culture media at a final concentration of 2 mM to increase neuron viability yet may not be enough or may only be one of the factors associated with both viability and plating density. Our improved protocol increases the neuron enrichment, as shown by qRT-PCR. However, other neural cells in culture, even if in low abundance, may also influence the density associated effects. Astrocytes release neurotrophic factors, such as BDNF, that augment neuron survival and function^82^. Endothelial cells may also be present in our culture, as we do not remove them with our protocol. These cells can secrete several factors, including GDNF, that can increase neuron survival and neurite growth^83,84^. While further elucidation is required to determine which organic molecules are released and by which cells, our data identified an appropriate cell density that allows to screen for compounds modulating neuron survival and neurite growth.

Laminin is a glycoprotein that is part of the extracellular matrix and is involved in cell differentiation, attachment, and growth^85^, partially through its interaction with integrins^86^. Laminin is commonly used as a precoating substrate to assist in the attachment of neurons and can increase the survival and growth of human pluripotent stem cell-derived neurons^87^. The addition of an extra layer of laminin coating on top of the PDL substrate further enhances attachment, survival, and growth of neural precursor cells^88^. Therefore, the addition of a laminin coating was tested on adult neurons with and without B27^+^ to assess its effects on cell attachment and growth. To our surprise, laminin did not impact neurite morphology, or the number of valid neurons and effects were not masked by B27^+^ supplementation. Therefore, the isolated adult cortical neurons do not seem receptive to laminin. We specifically used mouse derived laminin for this experiment to reduce the potential for inflammatory reactions and nonhomology between other species. We hypothesize that adult neurons have a reduced number of integrin receptors, or that downstream pathways are activated to a lesser extent relative to younger neurons resulting in reduced response to laminin coatings. Dorsal root ganglia neurons from P0 rats are able to increase integrin expression in culture while adult neurons are uncapable of this adaptation resulting in reduced survival and neurite outgrowth^89^. Transgenic expression of integrins in adult neurons restores their neurite regenerating capacity^90^. Therefore, integrin-dependent substrates, such as laminin, are not necessary to culture adult neurons for short period of time. This observation is in fact of high interest, as it reduces both the cost and time of screening, while minimizing the number of confounding variables. Indeed, not using laminin coatings will allow for screening molecules activating the downstream molecular signaling pathways without potential confounding factors.

The dissociation of neurons from the extracellular matrix is an essential part of neuron isolation. Traditional methods include incubating the brain tissue in digestive enzymes^91^, digesting through mechanical means^92^, or both^93^. To maximize neuronal yield, we opted for using both methods. Papain, which has been used in previous studies to dissociate neurons^94^, produced the highest number of valid neurons when used at concentrations ≥ 0.3 mg/mL. It is unknown if papain increases the enzymatic dissociation of neurons from the extracellular matrix or if it is gentler on adult neurons resulting in higher rates of survival compared to other dissociation enzymes. Different incubation times, temperatures, and mechanical dissociation methods were also tested to maximize the number of valid neurons. Lower temperatures and incubation times from Miltenyi’s original protocol^95^ were assessed to determine if milder conditions would increase neuron viability and survival. With the use of papain specifically, the reduction in temperature and time yielded significantly less valid neurons hypothesized to be from reduced enzyme efficiency as it is unconceivable how decreasing the temperature and incubation period would reduce the viability of neurons. A very large improvement to traditional methods was achieved with the gentleMACS™ Octo Dissociator with Heaters even when comparing to other methods of agitation. When no agitation method was used, the number of valid neurons were further reduced (data not included). It is unknown why the agitation induced by the gentleMACS™ Octo Dissociator with Heaters cannot be easily replicated using other methods. It may be providing the most effective amount of agitation while other methods may be over or under agitating the brain tissue. Some dissociation protocols require trituration, which are prone to bring great batch to batch variability^96^. Our protocol aims to maximize rigor and reducibility to be more sensitive to effect sizes. Through extensive testing, Miltenyi’s original dissociation method is the best tested method to increase the number of valid adult neurons. Figure 2 demonstrates the effects the dissociation enzymes have on the viability and growth capacity of neurons. The images also demonstrate that papain increases neuronal purity. RT-qPCR based RNA expression analysis confirmed the modified protocol (using papain instead of P&A) has higher neuronal yield and purity relative to Miltenyi’s original protocol which concurs with the images; automated morphology analysis only confirmed greater number of valid neurons as the other variables were not analyzed (Figure 2A-C). However, we did not conduct RNA expression analysis directly comparing the effects of substituting P&A with papain on neuronal yield and purity. Further experimentation is required to elucidate the effects of various dissociation enzymes on neuron health.

Many potent compounds are hydrophobic, and require the use of special solvents, which can be harmful to neurons. DMSO is a class 3 solvent (Food and Drug Administration) used to increase the solubility of hydrophobic compounds^61^. Therefore, many high content screens utilize DMSO to dissolve and prevent precipitation of hydrophobic compounds in cell culture media^62,63^. The issue of using DMSO as a vehicle is its toxicity to neurons and can alter their properties. A brief treatment of 0.05% DMSO induces neurophysiological changes to both hippocampal and cortical neurons^97^. DMSO concentrations ≥ 0.5% induce neurite retraction of primary embryonic neurons^98^. The effects of DMSO on adult cortical neurons have not need assessed before, and it is necessary to determine the best concentration minimizing the impact of DMSO on neurons while being high enough to allow the solubilization of most compounds tested in future screens. Based on our findings, it is recommended that the use of DMSO in high content screenings in adult neurons follow 2 criteria: 1) The use of DMSO is minimized and is ≤ 0.05% of the culture media; 2) The concentration of DMSO is kept consistent between all cohorts to reduce the confounding effect of DMSO on the analysis. The mechanism of such outcomes has yet to be elucidated. One possibility is the reduction of ERK phosphorylation. Indeed, phosphorylation of ERK, induced by alpha-lipoic acid in mouse neuroblastoma N2a cells, promotes neurite outgrowth^99^, while DMSO has been shown to reduce ERK phosphorylation in blood cells^100^. Therefore, DMSO may be reducing the neurite outgrowth potential of neurons by modulating the ERK pathway. DMSO also reduces the number of valid neurons (total neurite outgrowth of ≥10 μm) in a dose-dependent manner which is associated with a reduction in neurite outgrowth (Figure 6). Therefore, it is plausible the reduction in neurite outgrowth is merely from the reduction in neuron survival.

Due to the vast range of responses the general population has towards a given therapeutic^101,102^, it is vital to consider the age and sex factor in the development of new therapeutics. The cells isolated from that demographic are expected to respond differently as well. Here, we determined the demographic specific effects of RO48 and 7-epi Paclitaxel in our targeted screen to demonstrate the necessity for conducting demographic specific screens in neurons to ensure the compounds influence the target audience. Our *in vitro* cell cultures have identical conditions between age groups and are from very genetically similar mice. Thus, cell cultures allow us to determine how the neuron’s sex chromosomes (sex difference) or epigenetic profile (age-dependent effect) impact neuronal phenotype without known confounding variables. Through our specific processing and culturing methodology, we discovered both sex and age differences in response to exposure to different drugs.

We demonstrated that RO48 induces both sex- and age-dependent effects on neuron morphology and viability. RO48 activates mTORC1/2 and PI3K and decreases the phosphorylation of S6-926^64^ and S6K1 which are negative regulators of neurite growth^103^. Overall, RO48 present positive effects regardless of sex and age. RO48 significantly increases the average of neurite length in young adult neurons and increases the total neurite outgrowth only in older females. RO48 is very effective at increasing the number of valid neurons, which is expected since both mTOR and S6K1 increase neuron survival^104^. Intriguingly, the potency of RO48 to increase the number of valid neurons differs significantly with age and sex. One explanation for the smaller effect of RO46 at increasing the number of valid neurons with age is the age-dependent reduction in mitochondrial function. Indeed, S6K1 enhances mitochondrial ATP production^105^ which in important for axon growth and cell survival and could be a mechanism of action of RO48. However, we recently demonstrated that aging reduces neuronal mitochondrial capacity and efficiency^21^. Therefore, it is possible that the increase in ATP production in presence of RO48 is not as potent in older neurons, leading to a reduced activity of RO48. PI3K may also be involved in the age-dependent effects. PI3K decreases as cells age^106^. PI3K reduces neurite branch formation^107^ which would lead to an increase in the average neurite length. Since only the younger cohorts have increased neurite length in response to RO48, there may be an age-dependent response to PI3K modulation or age-dependent ability to express PI3K resulting in the significant differences in average neurite length in response to RO48. The single instance RO48 was not beneficial was in the average neurite length on middle-aged female neurons. There is a decrease in the ratio of phosphorylated mTOR in middle-aged naked mole rats compared to their young adult counterparts^108^ which is part of the mTOR activation process^109^. The mTOR pathway is an essential part of neurite formation^110–112^. Since there was a significantly greater increase in total neurite outgrowth in the older cohort in response to RO48 relative to their younger counterparts, and a greater increase in average neurite length in younger cohorts relative to the middle-aged female cohort, it may be plausible that mTOR phosphorylation induces neurite growth in all cohorts in a similar manner. The differences between age groups may be due to the management of neurite elongation. Indeed, we have shown at 7 DIV, neurons from aged mice have more branching^21^. This may be due to neurons from younger mice better consolidating and focusing neurite growth into few branches for the objective of producing axons. Nonetheless, direct experimentation is required to elucidate the pathways responsible these differences.

The sex differences of RO48 might be due to the activation of sex-dependent pathways. PI3K inhibition increases hepatic GSH content, antioxidant genes, and catalase in males but not in females^113^. The increase of catalase activity is correlated with reduction in neurite outgrowth^114^, therefore, one intriguing possibility is that the influence of PI3K on neurite outgrowth is sex-dependent.. Another possibility is the S6 phosphorylation rate difference. Indeed, female mice have significantly higher rates of S6 phosphorylation in both the liver and heart^115^, and therefore, may be more susceptible to the reduction in S6 phosphorylation induced by RO48^64^. This remains to be determined in cortical neurons. Inhibition of mTOR only increases brain proteasome activity in females^116^. Proteasome inhibition is a potential therapeutic option for increasing neurite outgrowth^117^, therefore, sex differences in neurite outgrowth through the mTOR-proteasome pathway would not surprising.

7-epi Paclitaxel is an FDA-approved drug for use in patients with ovarian cancer^65^. 7-epi Paclitaxel stabilizes microtubule bundles, impairs organelle transport^66^, and induces peripheral neuropathy through the CXCR1/2 pathway^67^. 7-epi Paclitaxel improved the average neurite length of all 3 demographics tested yet was toxic only to young adults demonstrated by reduced neurite outgrowth and number of valid neurons. However, severe neurotoxicity occurs sooner and more frequently in older metastatic breast cancer patients treated with 7-epi Paclitaxel compared to younger patients^118^. This discrepancy may be due to older patients generally being more vulnerable to neurotoxicity regardless of the treatment^119,120^. Conceivably, the increased 7-epi Paclitaxel-induced toxicity in older patients may be due to pharmacokinetic changes in aged patients that lead to prolonged drug activity and toxicity and increased drug sensitivity^121^. Therefore, it is plausible 7-epi Paclitaxel is less toxic to older neurons yet still more toxic to older patients. Interestingly, the CXCR1/2 expression is increased with age in rat cortical neurons^122^. The lower baseline of CXCR1/2 levels in younger neurons may explain the increased neurotoxicity relative to middle-aged neurons. Further direct experimentation is required to elucidate the age-dependent neurotoxicity of 7-epi Paclitaxel.

## Conclusion

Screening for compounds directly in adult cortical neurons has been so far unrealistic. Here, we established a new protocol that allows for screenings to be conducted directly in primary mouse adult cortical neurons. We conducted targeted screenings and demonstrated both age- and sex-dependent effects of multiple compounds. We established that: 1) dosage can intrinsically be sex-dependent; 2) screening in younger demographics will cause premature dismissal of compounds beneficial to older demographics; and 3) aging alters neuronal characteristics and therefore, must be considered in future screenings, especially for age-associated neurological disorders. This novel methodology is expected to strengthen the drug discovery process for neurological disorders and neurotraumatic injuries by providing more relevant *in vitro* data increasing the likelihood of preclinical and clinical success.

## Author Contributions

A.S. and C.G.G conceptualized, developed the methodology, designed the experiments, and wrote the manuscript. A.S. conducted all experiments. C.G.G. provided funding. I.R. provided equipment and resources to perform experiments. All authors have read and agreed to the published version of the manuscript.

## Acknowledgments

This research was funded by Texas A&M start-up funds to C.G.G. We thank Vance Lemmon, John Bixby, and Hassan Al-Ali for providing the RO48 compound and for their feedback on the methodology, analysis, and the overall manuscript. We thank Dwight Baker for providing consultation on assay development, and Jim Sacchettini for the use of the TAMU High Throughput Screening (HTS) Drug Screening Lab. We thank Graphit Science & Art for preparing the illustrations used in the figures.

